# An evolutionarily young defense metabolite influences the root growth of plants via the ancient TOR signaling pathway

**DOI:** 10.1101/150730

**Authors:** F.G. Malinovsky, M-L.F. Thomsen, S.J. Nintemann, L.M. Jagd, B. Bourgine, M. Burow, D. J. Kliebenstein

## Abstract

To optimize fitness a plant should monitor its metabolism to appropriately control growth and defense. Primary metabolism can be measured by the universally conserved TOR (Target of Rapamycin) pathway to balance growth and development with the available energy and nutrients. Recent work suggests that plants may measure defense metabolites to potentially provide a strategy ensuring fast reallocation of resources to coordinate plant growth and defense. There is little understanding of mechanisms enabling defense metabolite signaling. To identify mechanisms of defense metabolite signaling, we used glucosinolates, an important class of plant defense metabolites. We report novel signaling properties specific to one distinct glucosinolate, 3- hydroxypropyl glucosinolate across plants and fungi. This defense metabolite, or derived compounds, reversibly inhibits root growth and development. 3-hydroxypropyl glucosinolate signaling functions via genes in the ancient TOR pathway. Thus, plants might link evolutionarily new defense metabolites to ancient signaling pathways to optimize energy allocation.

## Introduction

Herbivory, pathogen attacks and weather fluctuations are just some of the factors that constantly fluctuate within a plants environment. To optimize fitness under this wide range of conditions, plants utilize numerous internal and external signals and associated signaling networks to plastically control metabolism and development (2-4). This metabolic and developmental plasticity begins at seed germination, where early seedling growth is maintained by heterotrophic metabolism relying solely on nutrients and energy stored in the seed including the embryo. Upon reaching light, the seedling transitions to autotrophy by shifting metabolism to initiate photosynthesis and alters development to maximize photosynthetic capacity (4, 5). Until light is available, it is vital for the plant to prioritize usage from the maternal energy pool, to ensure the shoot will breach the soil before resources are depleted. Because the time to obtaining light is unpredictable, seedlings that had the ability to to measure and accordingly adjust their own metabolism would likely enjoy a selective advantage. In this model, energy availability would an essential cue controlling growth throughout a plant’s life and not solely at early life-stages. On a nearly continuous basis, photo-assimilates, such as glucose and sucrose, are monitored and their internal levels used to determine the growth potential by partitioning just the right amount of sugars between immediate use and storage (3).

Illustrating the key nature of metabolite measurement within plants is that glucose, is measured by two separate kinase systems that are oppositely repressed and activated to determine the potential growth capacity, SnRKs1 (sucrose non-fermenting 1 (SNF1)-related protein kinases 1) and the Target of Rapamycin (TOR) kinase (6). SnRKs1s are evolutionarily conserved kinases that are activated when sugars are limiting (7). *Arabidopsis thaliana* (Arabidopsis) has two catalytic SnRK1-subunits, KIN10 and KIN11 (SNF kinase homolog 10 and 11), that activate vast transcriptional responses to repress energy-consuming processes and promote catabolism (8-10). This leads to enhanced survival during periods of energy starvation. Oppositely, the TOR kinase is a central developmental regulator, whose sugar-dependent activity controls a myriad of developmental processes including cell growth, cell-cycle, and cell-wall processes. The TOR pathway functions to modulate growth and metabolism by altering transcription, translation, primary and secondary metabolism, as well as autophagy (6, 11). The TOR kinase primarily functions in meristematic regions where it promotes meristem proliferation. Within these cell types, TOR measures the sugar content and if the tissue is low in sugar, TOR halts growth, even overruling hormone signals that would otherwise stimulate growth (12). In plants, TOR functions within a conserved complex that includes RAPTOR (regulatory-associated protein of TOR) and LST8 (lethal with sec-13 protein 8) (2). RAPTOR likely functions as an essential substrate-recruiting scaffold enabling TOR substrate phosphorylation (4), and LST8 is a seven WD40 repeats protein with unclear function (13). TOR complex (TORC) activity is positively linked with growth (4) as mutants in any component lead to qualitative or quantitative defects in growth and development and even embryo arrest in strong loss-of-function alleles (14, 15). Although the energy sensory kinases KIN10/11 and TOR sense opposite energy levels, they govern partially overlapping transcriptional networks, which are intimately connected to glucose-derived energy and metabolite signaling (6, 10). Having two systems to independently sense sugar shows the importance of measuring internal metabolism. A key pathway controlled by TOR in all eukaryotes is autophagy (16, 17). In non-stressed conditions, continuous autophagy allows the removal of unwanted cell components like damaged, aggregated or misfolded proteins by vacuolar/lysosomal degradation (18). Under low energy conditions, TORC inhibition leads to an induction of autophagy to free up energy and building blocks, through degradation of cytosolic macromolecules and organelles (17). Autophagy-mediated degradation is facilitated by formation of autophagosomes; double membrane structures that enclose cytoplasmic cargo, and delivers it to the vacuole (17, 19-21).

In nature, plant plasticity is not only limited to responding to the internal energy status, but to an array of external environmental inputs. The multitude of abiotic and biotic factors that plants continuously face often require choices between contradictory responses, that requires integrating numerous signals across an array of regulatory levels to create the proper answer. For example; plant defense against biotic organisms requires coordination of metabolic flux to defense and development while in continuous interaction with another organisms and the potential for interaction with other organisms. A proper defense response is vital for the plant as a metabolic defense response to one organism can impart an ecological cost by making the plant more sensitive to a different organism (22). Therefore, a plant must choose the most appropriate defense response for each situation to optimize its fitness and properly coordinate its defense response with growth and development. A key defense mechanism intricately coordinated across development is the synthesis of specific bioactive metabolites that are often produced in discrete tissues at specific times. A current model is that developmental decisions hierarchically regulate defense metabolism with little to no feed-back from defense metabolism to development. However, work on the glucosinolate and phenolic pathways is beginning to suggest that defense metabolites can equally modulate development (23-28), thus suggesting that development and defense metabolism can directly cross-talk.

To asses if and how defense metabolites can signal developmental changes, we chose to investigate the glucosinolate (GSL) defense metabolites. The evolution of the core of GSL biosynthesis is relatively young, and specifically modified GSL structures are even more recent (29). There are >120 known GSL structures limited to plants from the *Brassicales* order and some *Euphorbiaceae* family members, with Arabidopsis containing at least 40 structures (29). GSLs are amino acid derived defense metabolites that, after conversion to an array of bioactive compounds, provide resistance against a broad suite of biotic attackers (29-31). GSLs not only exhibit a wide structural diversity, but their composition varies depending on environmental stimuli, developmental stage and even across tissues. This, combined with the information-rich side chain, makes GSLs not only an adaptable defense system, but also prime candidates for having distinctive signaling functions. Previous work has suggested that there may be multiple signaling roles within the GSLs (27, 28, 32-34). Tryptophan-derived indole GSLs or their breakdown products alter defense responses to non-host pathogens illustrated by biosynthetic mutants devoid of indole GSLs being unable to deposit PAMP-induced callose in cell walls via an unknown signal and pathway (32, 34). Similarly, indole GSL activation products have the ability to directly alter auxin perception by interacting with the TIR1 auxin receptor (23). In contrast to indole GSLs, mutants in aliphatic GSL accumulation alter flowering time and circadian clock oscillations (35, 36). The aliphatic GSL activation product, allyl isothiocyanate can induce stomatal closure but it is unknown if this is specific to allyl GSL or a broader GSL property (33, 37). Allyl GSL (other names are 2-propenyl GSL and sinigrin) can also alter plant biomass and metabolism in Arabidopsis (27, 28). While these studies have provided hints that the GSL may have signaling potential, there is little understanding of the underlying mechanism or the structural specificity of the signal. To explore whether built-in signaling properties are a common attribute of GSLs, we screened for altered plant growth and development in the presence of specific purified aliphatic GSLs. In particular, we were interested in identifying candidate signals whose activity could ensure fast repartitioning of resources between development and defense. Here we present a novel signaling capacity specific to the aliphatic 3-hydroxypropyl glucosinolate (3OHP GSL). Our results suggest that 3OHP GSL signaling involves the universally conserved TOR pathway for growth and development, as mutants in TORC and autophagy pathways alter responsiveness to 3OHP GSL application. As such, our findings point towards that evolutionarily young defense metabolites can signal plastic changes in plant development by interacting with highly conserved signaling pathways.

## Results

### 3OHP GSL inhibits root growth in Arabidopsis

We reasoned that if a GSL can prompt changes in plant growth it is an indication of an inherent signaling capacity. Using purified compounds, we screened for endogenous signaling properties among short-chain methionine-derived aliphatic GSLs by testing their ability to induce visual phenotypic responses in Arabidopsis seedlings. We found that 3OHP GSL causes root meristem inhibition, at concentrations down to 1μM (Fig. 1A). The observed response is concentration-dependent (Figure 1A-B). All Arabidopsis accessions accumulate 3OHP GSL in the seeds, and this pool is maintained at early seedling stages (38-40). Previous research has shown that GSLs in the seed are primarily deposited in the embryo, accumulating to about 3μmol/g suggesting that we are working with concentrations within the endogenous physiological range (41, 42).

**Figure 1.**
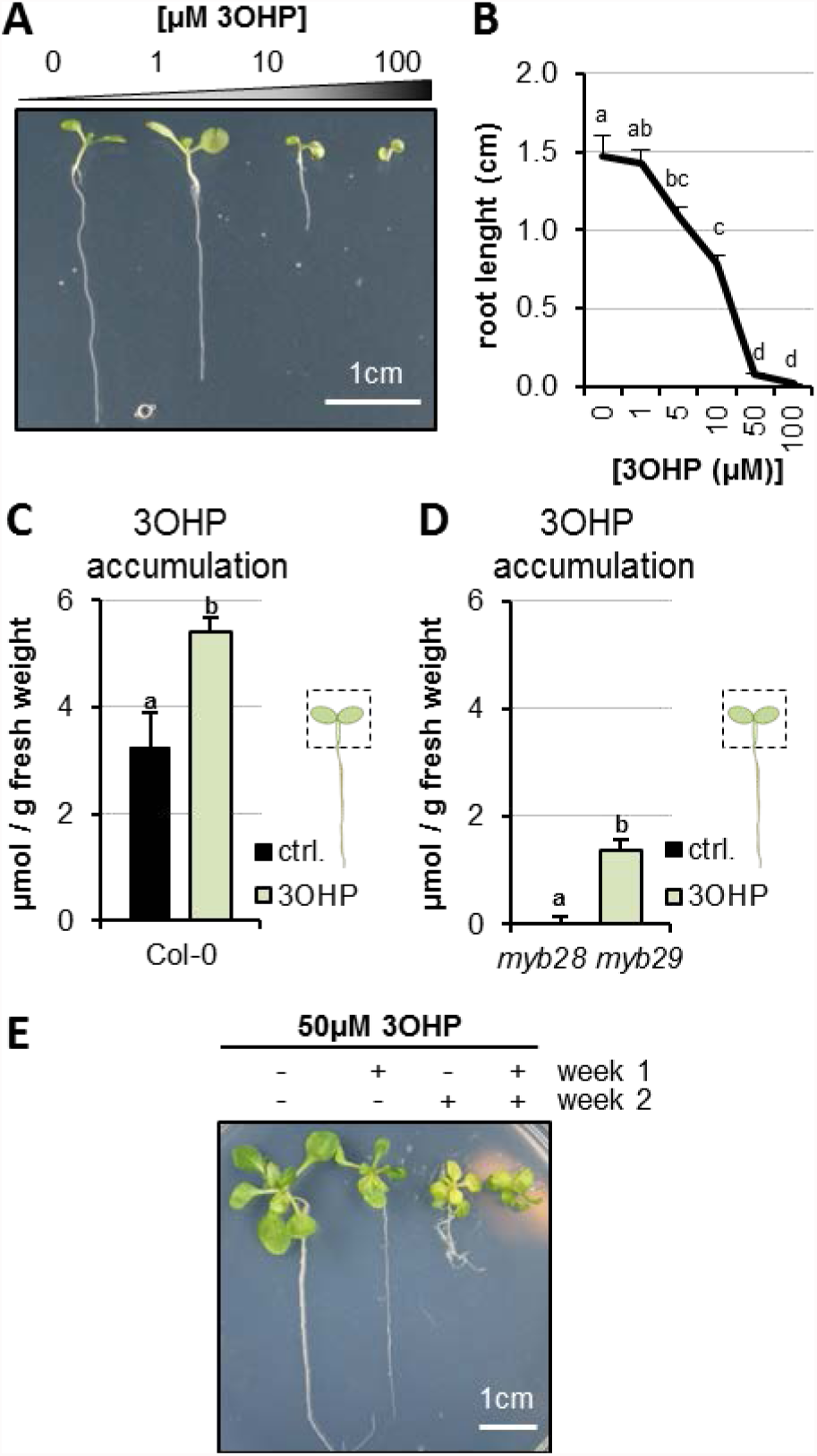
3OHP reversibly inhibits root growth. **A** 7-d-old seedlings grown on MS medium supplemented with a concentration gradient of 3OHP. **B** Quantification of root lengths of 7-d-old. Results are averages ± SE (n = 3-7; P < 0.001). **C** Accumulation of 3OHP in shoots/areal tissue of 10-d-old Col-0 wildtype seedlings grown on MS medium supplemented with 5μM 3OHP. Results are least squared means ± SE over three__independent experimental replicates with each experiment having an average eleven replicates of each condition (n = 31-33; ANOVA P_Treat_ < 0.001). **D** Accumulation of 3OHP in shoots of 10-d-old *myb28 myb29* seedlings (aliphatic GSL-free) grown on MS medium supplemented with 5μM 3OHP. Results are least squared means ± SE over two independent experimental replicates with each experiment having an average of four independent biological replicates of each condition (n = 8-14; ANOVA P_Treat_ < 0.001). **E** 14-d-old seedlings grown for 1 week with or without 3OHP as indicated. After one week of development, the plants were moved to the respective conditions showed in week 2.

We tested how exogenous 3OHP GSL exposure to the roots alters 3OHP GSL accumulation within the shoot, and how this compares to endogenously synthesized 3OHP GSL levels. We grew the Col-0 reference accession and the *myb28-1 myb29-1* mutant that is devoid of endogenous aliphatic GSLs in the presence and absence of exogenous 3OHP GSL. At day 10, the foliar 3OHP GSL levels were analyzed. Col-0 without treatment had average foliar levels of 3OHP GSL of 3.2 μmol/g and grown on media containing 5μM, 3OHP GSL contributed an additional 2.2 μmol/g raising the total 3OHP GSL to 5.4 μmol/g (Figure 1C). The *myb28-1 myb29-1* mutant had no measurable 3OHP GSL on the control plates and accumulated ~1.4μmol/g upon treatment (Figure 1D). In agreement with the lower foliar 3OHP GSL accumulation in the *myb28-1 myb29-1* mutant background, this double mutant had a lower root growth response to exogenous 3OHP GSL (Figure. 1 –figure supplement 1). Importantly, this confirms that the level of 3OHP GSL application is within the physiological range.

We then tested if 3OHP GSL or potential activation products inhibit root growth because of cell death or toxicity. The first evidence against toxicity came from the observation that even during prolonged exposures, up to 14 days of length, Col-0 seedlings continued being vital and green (Figure 1E). If there was toxicity the seedlings would be expected to senesce and die. We next tested if the strong root growth inhibition by 50μM 3OHP GSL is reversible. Importantly, root inhibition is reversible, as the 3OHP GSL-mediated root stunting could be switched on and off by transfer between control media and media containing 3OHP GSL (Figure 1E). Based on the toxicity and GSL assays, we conclude that the 3OHP GSL treatments are at reasonable levels compared to normal Arabidopsis physiology, and that the phenotypic responses we observed were not caused by flooding the system with 3OHP GSL or toxicity.

### Root inhibition is specific to 3OHP GSL

To evaluate whether 3OHP GSL mediated root inhibition is a general GSL effect or if it is structurally specific to 3OHP GSL, we tested if aliphatic GSLs with similar side-chain lengths, but different chain modifications, would induce similar root growth effects. First, we assessed 3- methylsulfinyl-propyl (3MSP) GSL, the precursor of 3OHP GSL, and the alkenyl-modified three carbon glucosinolate allyl (Figure 2A). In contrast to 3OHP GSL, neither of these structurally related GSLs possessed similar root-inhibiting activities within the tested concentration range (Figure 2B-D). We also analyzed the potential root inhibition for the one carbon longer C4-GSLs 4- methylsulfinylbutyl (4MSB) and but-3-enyl (Figure 2E). Neither of these compounds could inhibit root growth at the tested concentrations (Figure 2F-H). There is no viable commercial, synthetic or natural source for the 4-hydroxybutyl GSL which prevented us from testing this compound. The fact that only 3OHP GSL inhibits root elongation suggests that the core GSL structure (comprised of a sulfate and thioglucose) does not cause the effect. Importantly, this indicates that 3OHP GSL root inhibition is not a generic result of providing extra sulfur or glucose from the GSL core structure to the plant, as these compounds would be equally contributed by the other GSLs. Furthermore, the results confirm that there is no general toxic activity when applying GSLs to Arabidopsis. This evidence argues that the 3OHP GSL root inhibition effect links to the specific 3OHP side chain structure, indicating the presence of a specific molecular target mediating the root inhibition response.

**Figure 2.**
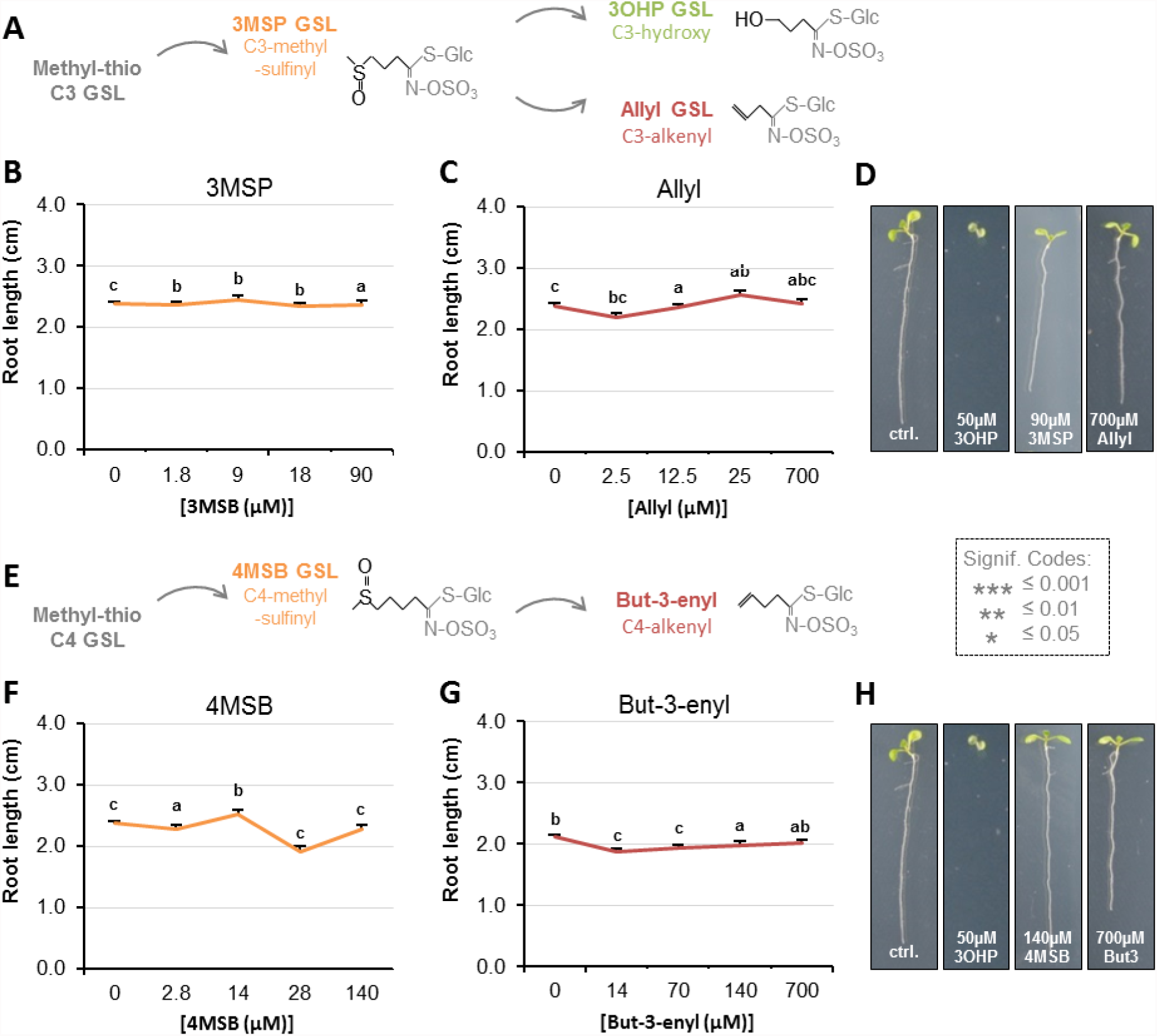
Root growth is not inhibited by all aliphatic GSLs. **A** The aliphatic glucosinolate biosynthetic pathway, from the C3 3-methyl-sulphinyl-propyl (3MSP) to the secondary modified 3-hydroxyl-propyl (3OHP) and 2-propenyl (allyl/sinigrin). **B-C** Root lengths of 7- d-old Col-0 wildtype seedlings grown on MS medium supplemented with a concentration gradient of the indicated aliphatic C3-GSL. The left most point in each plot shows the root length grown in the absence of the specific GSL treatment. Results are least squared means ± SE over four independent experimental replicates with each experiment having an average of 21 replicates per condition (n_3MSP_=59-153; n_Allyl_=52-153). Significance was determined via two-way ANOVA combining all experiments. **D** 7-d-old seedlings grown on MS medium with or without 50μM of the indicated GSL. **E** The aliphatic glucosinolate biosynthetic pathway from the C4 4-methyl-sulphinyl-butyl (4MSB) to But-3- enyl. **F-G** Root lengths of 7-d-old Col-0 wildtype seedlings grown on MS medium supplemented with a concentration gradient of the indicated aliphatic C4-GSL. The left most point in each plot shows the root length grown in the absence of the specific GSL treatment. Least squared means ± SE over four independent experimental replicates with each experiment having an average of 22 replicates condition (n_4MSB_=38-153; n_But-3-enyl_=68-164). Significance was determined via two-way ANOVA combining all experiments. **H** 7-d-old seedlings grown on MS medium with or without 50μM of the indicated GSL.

### 3OHP GSL responsiveness is wider spread in the plant kingdom than GSL biosynthesis

The evolution of the GSL defense system is a relatively young phylogenetic event that occurred within the last ~92 Ma and is largely limited to the Brassicales order (43). The aliphatic GSL pathway is younger still (~60 Ma) and is limited to the Brassicaceae family with the enzyme required for 3OHP GSL production, AOP3, being limited to *Arabidopsis thaliana* and *Arabidopsis lyrata* within the Arabidopsis lineage (43, 44). However, 3OHP GSL is also found in the vegetative tissue of the close relative *Olimarabidopsis pumila* (dwarf rocket) (45), and in seeds of more distant Brassicaceae family members such as the hawkweed-leaved treacle mustard (*Erysimum hieracifolium*), virginia stock (*Malcolmia maritima*), shepherd’s cress (*Teesdalia nudicaulis*), and alpine pennycress (*Thlaspi alpestre*) (46, 47). These species are evolutionarily isolated from each other, suggesting that they may have independently evolved the ability to make 3OHP GSL (44, 45, 48). As such, 3OHP GSL is an evolutionarily very young compound and we wanted to determine if the molecular pathway affected by 3OHP GSL is equally young, or whether 3OHP GSL affects an evolutionarily older, more conserved pathway.

First we tested for 3OHP GSL responsiveness in plant species belonging to the GSL-producing Brassicales order (Figure 3A). We found that 4 of the 5 tested Brassicales species responded to 5μM 3OHP GSL with root growth inhibition regardless of their ability to synthesize 3OHP GSL (Figure 3B-F). This suggests that responsiveness to 3OHP GSL application does not link to the ability to make 3OHP GSL. We expanded the survey by including plants within the eudicot lineage that do not have the biosynthetic capacity to produce any GSLs (Figure 3G) and found that 5μM 3OHP GSL can inhibit root growth in several of the non-Brassicales species tested (Figure 3H-L). The ability of 3OHP GSL to alter growth extended to *Saccharomyces cerevisae* where 3OHP GSL led to slower log phase growth than the untreated control (Figure 3 – Figure Supplement 1). Allyl GSL in the media had no effect on *S. cerevisae* growth showing that this was a 3OHP GSL mediated process (Figure 3 – Figure Supplement 1). The observation that 3OHP GSL responsiveness is evolutionarily older than the ability to synthesize 3OHP GSL suggests that the molecular target of 3OHP GSL, or derived compounds, must be present and highly conserved among these species. Similarly, if the signaling event occurs from a 3OHP GSL derivative, then the metabolic processes enabling the formation of this derivative are conserved beyond Brassicaceous plants.

**Figure 3.**
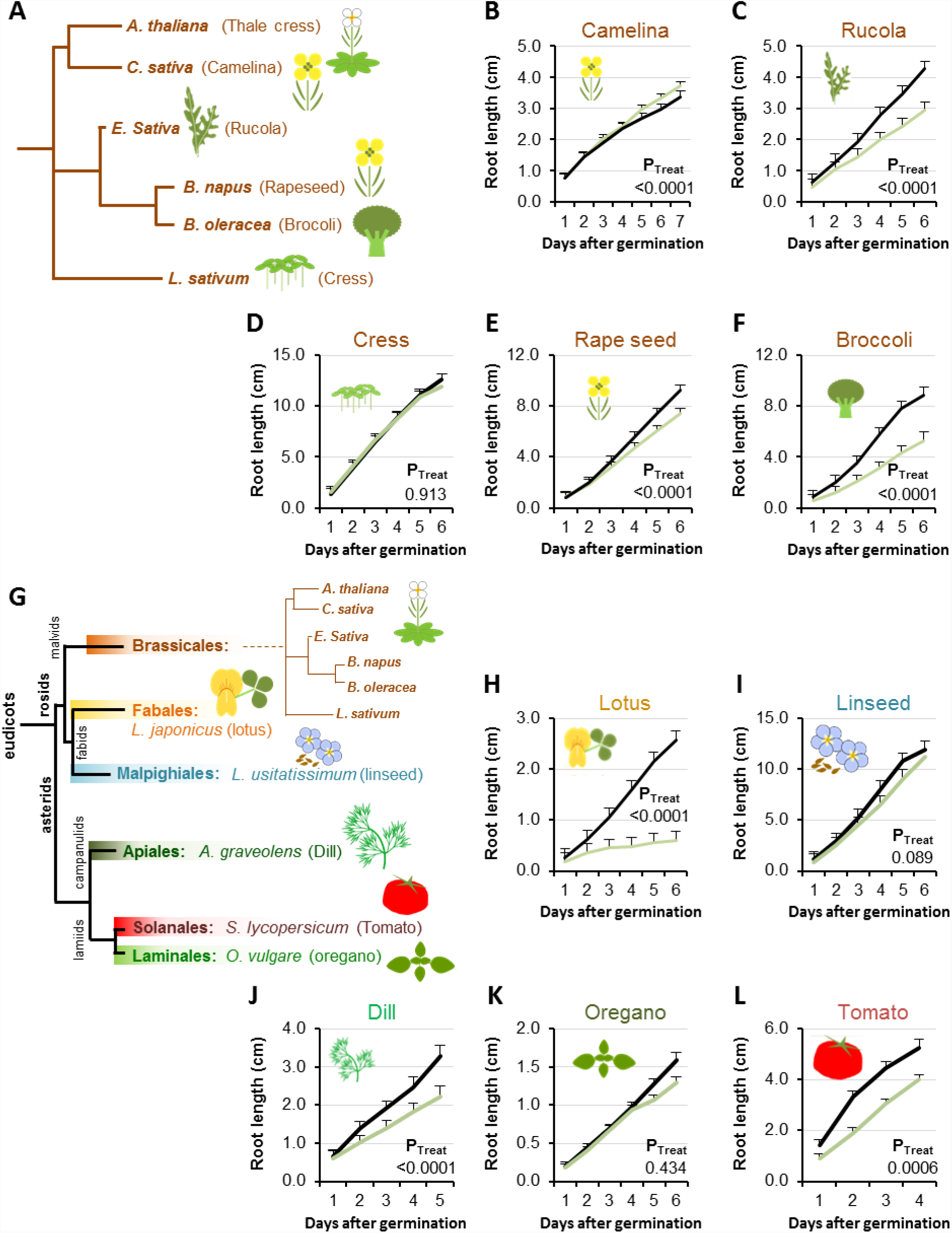
Conservation of 3OHP responsiveness suggests a evolutionally conserved target. **A** Stylized phylogeny showing the phylogenetic relationship of the selected plants from the Brassicales family, branch lengths are not drawn to scale. **B-F** plants from the Brassicales family, grown on MS medium supplemented with or without 5μM 3OHP. **G** Stylized phylogeny showing the phylogenetic relationship of all the selected crop and model plants, branch lengths are not drawn to scale. **H-L** Root growth of plants from diverse eudicot lineages, grown on MS medium supplemented with or without 5μM 3OHP. Results are least squared means ± SE for each species using the following number of experiments with the given biological replication. Camelina three independent experimental replicates (n_ctrl_=8 and n_3OHP_=12). Rucola three independent experimental replicates (n_ctrl_=17 and n_3OHP_=17. Cress; three independent experimental replicates (n_ctrl_=19 and n_3OHP_=18). Rape; seed four independent experimental replicates (n_ctrl_=14 and n_3OHP_=13). Broccoli; three independent experimental replicates (nctrl=10 and n_3OHP_=13). Lotus; three independent experimental replicates (n_ctrl_=10 and n_3OHP_=10). Linseed; three independent experimental replicates (n_ctrl_=11 and n_3OHP_=11). Dill; three independent experimental replicates (n_ctrl_=14 and n_3OHP_=13). Oregano; four independent experimental replicates (n_ctrl_=40 and n_3OHP_=39). Tomato; three independent experimental replicates (n_ctrl_=11 and n_3OHP_=15). A significant effect of treatment on the various species was tested by two-way ANOVA combining all the experimental replicates in a single model with treatment as a fixed effect and experiment as a random effect

### 3OHP reduces root meristem and elongation zone sizes

We hypothesized that 3OHP GSL application may alter root cellular development to create the altered root elongation phenotype. A reduction in root growth can be caused by inadequate cell division in the root meristematic zone or by limited cell elongation in the elongation zones (Figure 4A) (2). To investigate how 3OHP GSL affects the root cellular morphology, we used confocal microscopy of 4-d-old Arabidopsis seedlings grown vertically with or without 10μM 3OHP GSL. We used propidium iodide stain to visualize the cell walls of individual cells, manually counted the meristematic cells, and measured the distance to the point of first root hair emergence. Root meristems of 3OHP GSL treated seedlings were significantly reduced in cell number compared to untreated controls (Figure 4B-C). Moreover, we also observed a premature initiation of the differentiation zone, as the first root hairs were closer to the root tip upon 3OHP GSL treatment (Figure 4 D-E). In addition, we saw bulging and branching of the root hairs in 3OHP GSL treated roots (Figure 4F). There was no morphological evidence of cell death in any root supporting the argument that 3OHP GSL is not a toxin. These results indicate that 3OHP GSL leads to root growth inhibition by reducing the size of the meristematic zone within the developing Arabidopsis root.

**Figure 4.**
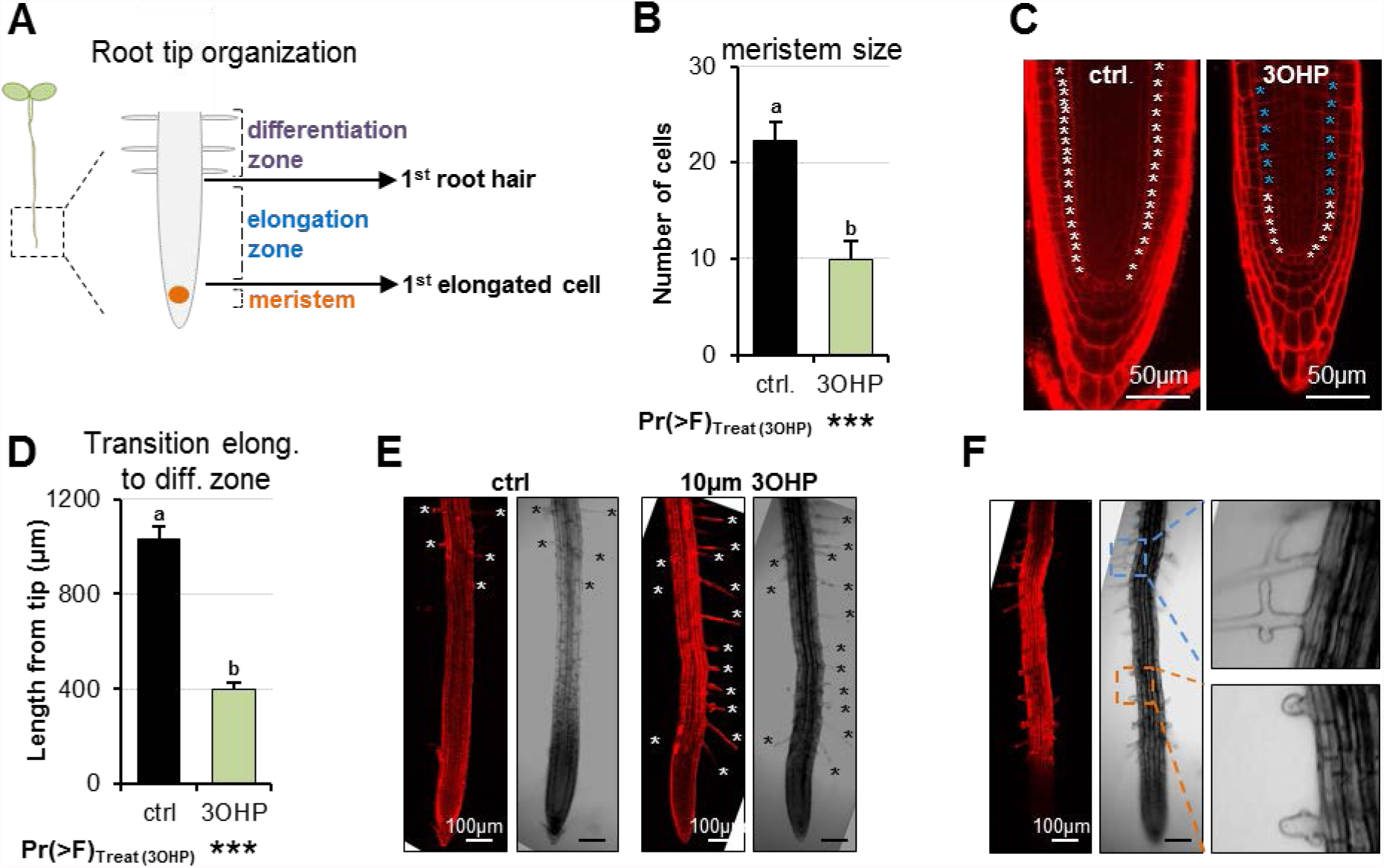
3OHP reduces root zone sizes. **A** Diagrammatic organization of a root tip; the meristem zone from the QC to the first cell elongation; the elongation zone ends when first root hair appears (1). **B** Meristem size of 4-d-old Arabidopsis seedlings grown on MS medium with sucrose ± 10μM 3OHP. Results are least squared means ± SE over three independent experimental replicates with each experiment having an average of three replicates per condition (n_ctrl_=6; n_3OHP_=9). Significance was tested via two-way ANOVA with treatment as a fixed effect and experiment as a random effect. **C** Confocal images of 4-d-old propidium iodide stained seedlings grown with and without 3OHP. Meristematic cells are marked with white asterisks, elongated cells with blue asterisks. **D** Appearance of first root hair; measured from the root tip on 4-d-old seedlings grown on MS medium with sucrose ± 10μM 3OHP. Results are least squared means ± SE over two independent experimental replicates with each experiment having an average of nine replicates per condition (n_ctrl_=17; n_3OHP_=20). Significance was tested via two-way ANOVA with treatment as a fixed effect and experiment as a random effect. **E** Confocal images of 4–d-old propidium iodide stained seedlings grown with and without 3OHP. Protruding root hairs are marked with white/black asterisks. **F** 3OHP induced root hair deformations, confocal images of 4–d-old propidium iodide stained seedlings grown with 3OHP.

### TORC-associated mutants alter 3OHP GSL responsiveness

The observed response to 3OHP GSL suggests that the target of this compound is evolutionarily conserved and alters root growth but does not affect the patterning of the root meristem. This indicates that key root development genes like SHR and SCR are not the targets as they affect meristem patterning (49, 50). Mutants in GSL biosynthetic genes can lead to auxin over-production phenotypes as indicated by the superroot (*SUR*) 1 and 2 loci (51, 52). However, the *SUR* genes are not evolutionarily conserved and 3OHP GSL does not create a superroot phenotype, showing that the genes are not the targets. A remaining conserved root regulator that does not alter meristem formation, but still alters root growth, is the TOR pathway (12). Thus, we proceeded to test if mutants in the TOR pathway alter sensitivity to 3OHP GSL. Because TORC activity is sugar responsive, we investigated whether 3OHP GSL application may alter the response to sugar in genotypes with altered TORC activity. We first used the TOR kinase overexpression line GK548 (TORox) because it was the only one of several published TOR overexpression lines (53) that behaved as a TOR overexpressor within our conditions (Figure 5 –figure supplement 1). The GK548 TORox line exhibits accelerated TORC signaling and consequently grows longer roots on media containing sucrose (Figure 5 and (53)). In addition, GK548 TORox meristems are harder to arrest (Figure 5B). Applying 3OHP GSL to the GK548 TORox line showed that this genotype had an elevated 3OHP GSL-mediated inhibition of meristem reactivation in comparison to the WT (Figure 5C-D). This suggests that TORC activity influences the response to 3OHP GSL (Figure 5E).

**Figure 5.**
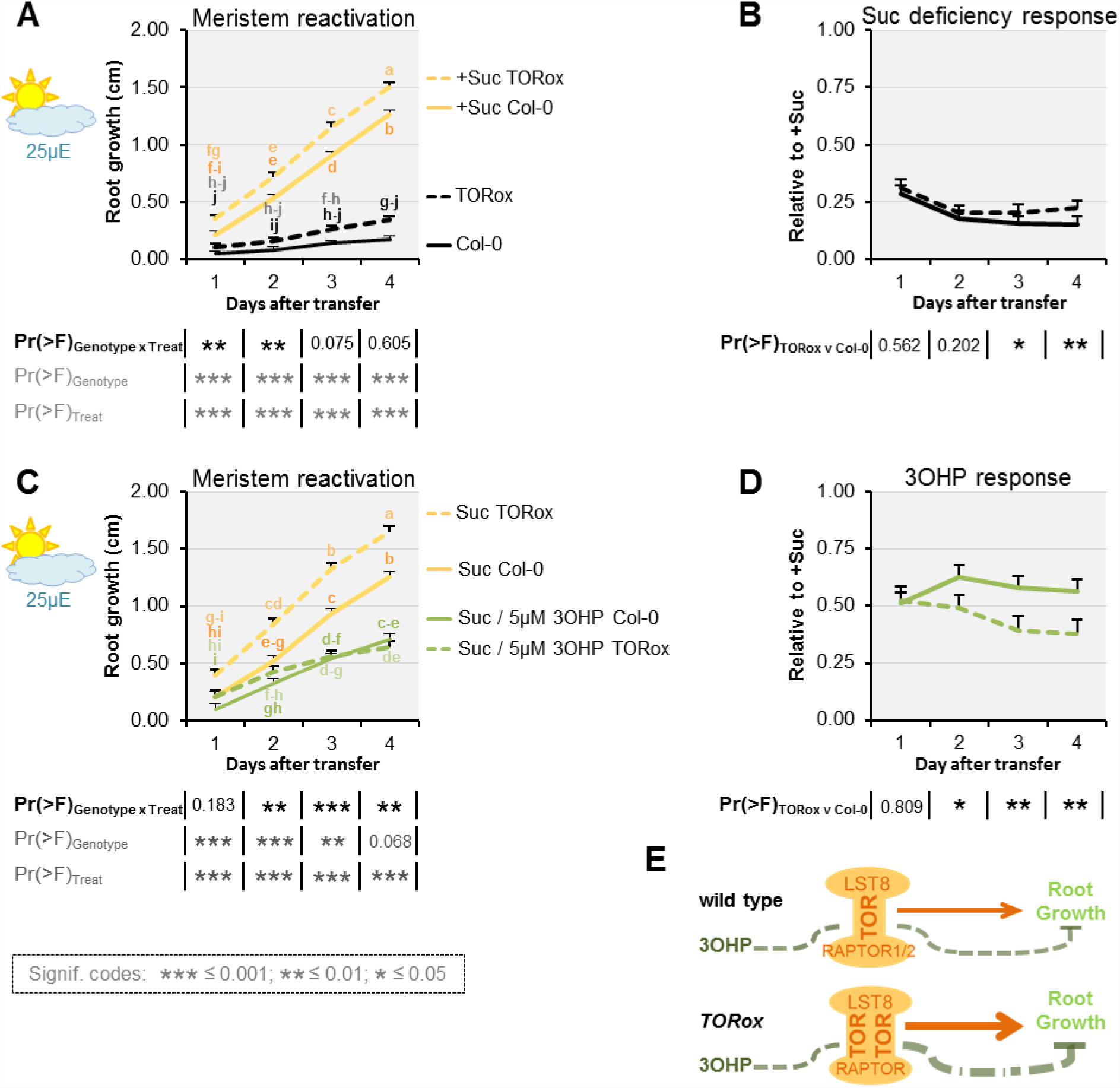
TOR over-activation amplifies 3OHP response. **A** Root growth for low light grown seedlings. The seedlings were grown on MS medium without sucrose for 3 days, then transferred to the indicated media (Suc; sucrose). Multi-factorial ANOVA was used to test the impact of Genotype (Col-0 v TORox), Treatment (Control v Sucrose) and their interaction on root length. All experiments were combined in the model and experiment treated as a random effect. The ANOVA results from each day are presented in the table. **B** The root lengths grown photo-constrained and without sucrose (from A) displayed at each time point as relative to the respective sucrose activated roots. Results least squared means ± SE over three independent experimental replicates with each experiment having an average of nine replicates per condition (n=26-30). Multi-factorial ANOVA was used to test the impact of Genotype (Col-0 v TORox), Treatment (Sucrose v Sucrose/3OHP) and their interaction on root length. All experiments were combined in the model and experiment treated as a random effect. The ANOVA results from each day are presented in the table. **C** Root growth for low light grown seedlings. The seedlings were grown on MS medium without sucrose for 3 days, then transferred to the indicated media. **D** Photo-constrained root lengths in response to sucrose and 3OHP (from A) displayed at each time point as relative to the respective sucrose activated roots. Results are least squared means ± SE over two independent experimental replicates with each experiment having an average of six replicates per condition (n=11-14). **E** Schematic model; over

We next investigated how genetically disrupting additional components of TORC affects 3OHP GSL responsiveness. In addition to the catalytic TOR kinase subunit, TORC consists of the substrate binding RAPTOR (2, 4), and LST8 (13) (Figure 5E). In Arabidopsis- there is one copy of *TOR*, and two copies of both *RAPTOR* and *LST8* (*RAPTOR1*/*RAPTOR2* and *LST8-1*/*LST8-2*) (13, 54).

RAPTOR1 and TOR null mutants are lethal as homozygotes (14, 15), and heterozygous *raptor1* mutants did not display a significant change in 3OHP GSL responsiveness (Figure. 5 –figure supplement 2). We therefore tested insertion mutants within the weaker homolog *RAPTOR2*, whose null mutant is viable, and in our conditions shows mildly reduced root length on sucrose-containing media (Figure 5 –figure supplement 3A and C). We found that, for two independent insertion lines *raptor2-1* (54, 55) and *raptor2-2* (54) (Figure 5 –figure supplement 3E), there was a statistically significant reduction in 3OHP GSL response (Figure 5 –figure supplement 3A-C). This supports the hypothesis that 3OHP GSL-associated signaling proceeds through TORC and that *RAPTOR2* may play a stronger role in 3OHP perception than *RAPTOR1*.

### 3OHP GSL treatment inhibits sugar responses

A key function of TORC activity is to control meristem cell division and this can be measured by meristem reactivation assays (12). Thus, to further test if TORC dependent responses are altered by 3OHP GSL, seedlings were germinated in sugar-free media and photosynthesis-constrained under low light conditions to induce root meristem arrest when the maternal glucose is depleted (three days after germination). The root meristems were reactivated by applying exogenous sucrose (Figure 6A-C). By treating arrested root meristems with sucrose alone or in combination with 3OHP GSL we found that 3OHP GSL could inhibit meristem reactivation of sugar-depleted and photosynthesis-constrained seedlings (Figure 6D-E). Further, this response was dependent upon the 3OHP GSL concentration utilized. A similar response was found when treating with a TOR inhibitor such as rapamycin (12), providing additional support to the hypothesis that 3OHP GSL may reduce root growth by altering TORC activity.

**Figure 6.**
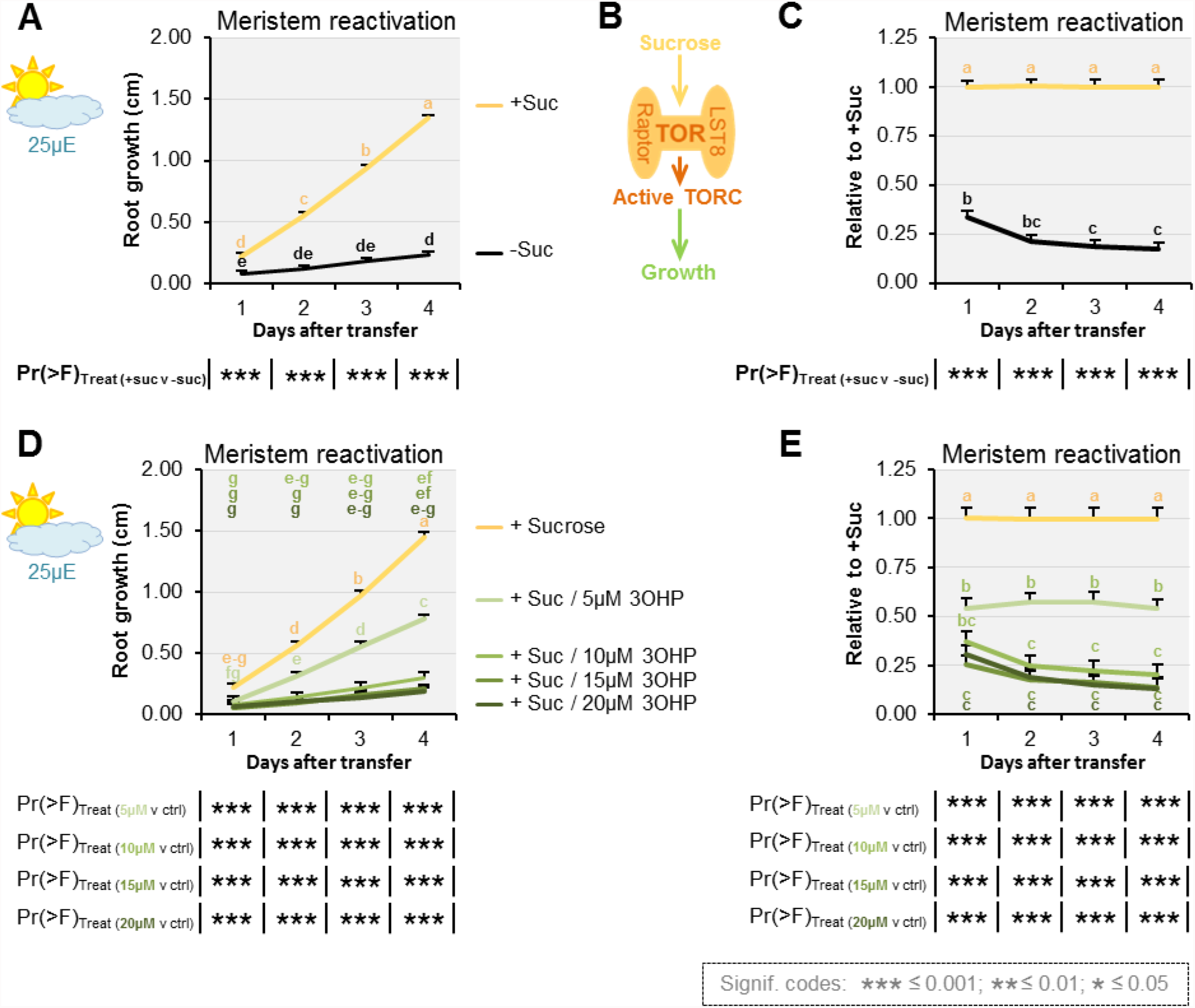
3OHP dampens sugar-mediated meristem activation. **A** Root growth for low light grown Col-0 wildtype seedlings. The seedlings were grown on MS medium without sucrose for 3 days, then transferred to the indicated media. Multi-factorial ANOVA was used to test the impact of Treatment on root length. All experiments were combined in the model and experiment treated as a random effect. The ANOVA results from each day are presented in the table. **B** Schematic model; sucrose activates the TOR complex (TORC), leading to growth. **C** The root lengths (from A) displayed at each time point as relative to sucrose activated roots. Results are least squared means ± SE over five independent experimental replicates with each experiment having an average of eight replicates per condition (n_-Suc_= 43; n_+Suc_=40). **D** Root growth for low light grown seedlings. The seedlings were grown on MS medium without sucrose for 3 days, then transferred to the indicated media. Multi-factorial ANOVA was used to test the impact of Treatment on root length. All experiments were combined in the model and experiment treated as a random effect. The ANOVA results from each day are presented in the table. **E** The root lengths (from D) displayed at each time point as relative to sucrose activated roots (ctrl.). Results are least squared means ± SE over two independent experimental replicates with each experiment having an average of seven replicates per condition (n=12-16).

### 3OHP GSL pharmacologically interacts with the TOR-inhibitor AZD-8055

To further examine the possibility that 3OHP GSL may be affecting the TOR pathway, we proceeded to compare the effect of 3OHP GSL to published chemical TOR inhibitors. The active site TOR inhibitors were originally developed for mammalian cells and inhibit root growth in various plant species (56). Similar to 3OHP GSL, the active-site TOR inhibitor AZD-8055 (AZD) induces a reversible concentration-dependent root meristem inhibition (56). By directly comparing 3OHP GSL treatment with known TOR chemical inhibitors in the same system, we can test for interactions between 3OHP GSL and the known TOR inhibitors. An interaction between 3OHP GSL application and a known TOR inhibitor, e.g. an antagonistic relationship, is an indication that the same target is affected. To assess whether interactions between 3OHP GSL and TOR signaling occur, we grew seedlings vertically on media with combinations of 3OHP GSL and AZD and root-phenotyped the plants to compare the effect on root morphology. This identified a significant antagonistic interaction between AZD and 3OHP GSL (3OHP x AZD), both in terms of root length response (Figure 7A) and in initiation of the differentiation zone (Figure 7B). This antagonistic interaction is also supported by the appearance of first root hair (Figure 7B), as the premature initiation of the differentiation zone in the presence of 10 μM 3OHP GSL did not change further upon co-treatment (Figure 7B). Moreover, there was a vast overlap in the phenotypic response to both compounds (Figure 7C-D); notably the closer initiation of the root differentiation zone to the root tip (Figure 7B-C) and the decreased cell elongation (Figure 7C, right panel). Together, this suggests that the TOR inhibitor AZD and 3OHP GSL have a target in the same signaling pathway as no additive effect is observed. Supporting this is the observation that 3OHP GSL treatment is phenotypically similar to a range of TOR active site inhibitors, as well as an inhibitor of S6K1 (one of the direct targets of TOR) (Figure 7 –figure supplement 1).

**Figure 7.**
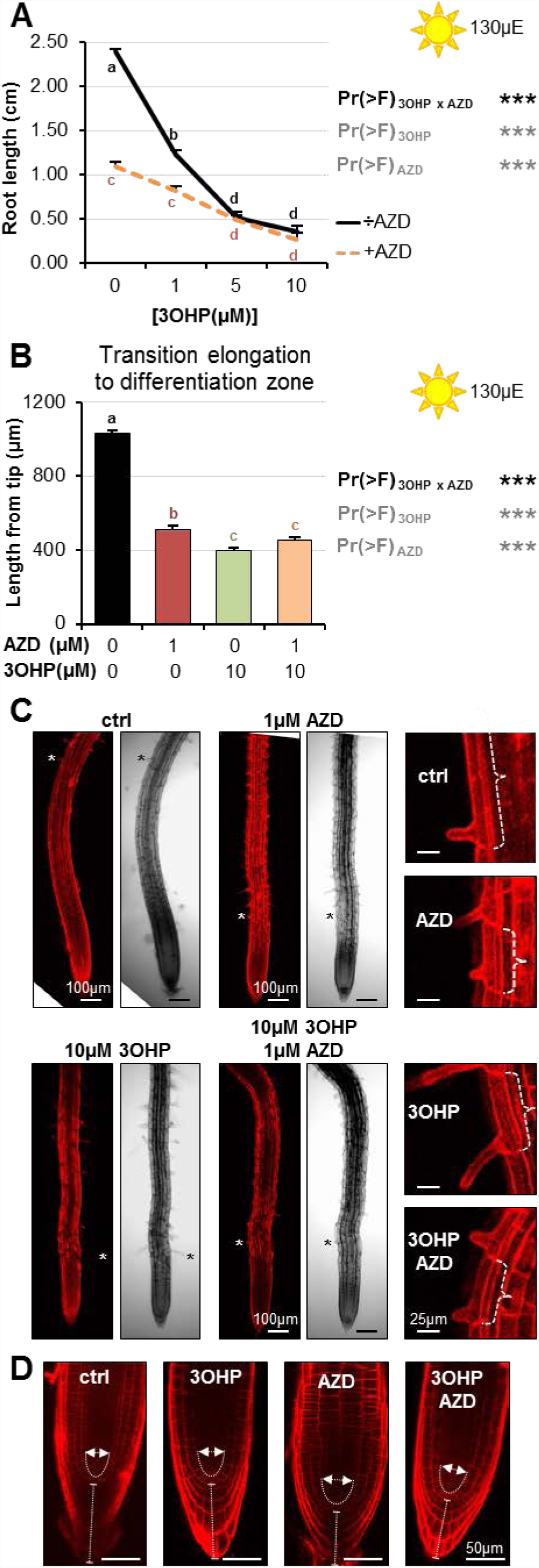
**A** Root lengths of 7-d-old Col-0 wildtype seedlings grown on MS medium with sucrose ± combinations of AZD and different concentrations of 3OHP. Results are least squared means ± SE over three independent experimental replicates with each experiment having an average of nine replicates per condition (n=18-58). Multi-factorial ANOVA was used to test the impact of the two treatments and their interaction on root length. All experiments were combined in the model and experiment treated as a random effect. The ANOVA results from each day are presented in the table. **B** Appearance of first root hair; measured from the root tip on 4–d-old seedlings grown on the indicated MS medium with sucrose. Results are least squared means ± SE over two independent experimental replicates with each experiment having an average of nine replicates per condition (n=17-20). Multi-factorial ANOVA was used to test the impact of the two treatments and their interaction on root length. All experiments were combined in the model and experiment treated as a random effect. The ANOVA results from each day are presented in the table. **C** Confocal images of 4- d-old propidium iodide stained seedlings. The first protruding root hairs are marked with white/black asterisks on the left panel. Right panel shows zooms of first root hair, cell size is indicated. **D** Confocal images of 4–d-old propidium iodide stained seedlings.

Interestingly, the short root hair phenotype induced by AZD showed a synergistic interaction between AZD and 3OHP GSL suggesting that they may target different components of the TORC pathway that interact (Figure 7C). Further, while there is strong phenotypic overlap between AZD and 3OHP GSL, there are also specific activities. AZD induced a rounding of the root tip (1), but co-treatment with 3OHP GSL restored a wildtype-like tip phenotype (Figure 7D). The lack of root rounding and root hair inhibition suggest that AZD and 3OHP GSL both target the TOR pathway, but at different positions. Alternatively, the 3OHP GSL may be a more specific TOR inhibitor and the additional AZD phenotypes could be caused by the ATP-competitive inhibitor having alternative targets in plants. Together, these results suggest that 3OHP GSL directly or indirectly targets the same molecular pathway as known TOR inhibitors (57).

### Blocking parts of the autophagy machinery affects 3OHP GSL associated signaling

Activation or repression of the TOR pathway leads to regulatory shifts in numerous downstream pathways (Figure 8E) (6, 11). For example, active TOR negatively regulates autophagy across eukaryotic species including Arabidopsis (16, 17). To test if pathways downstream of TORC are affected by, or involved in, 3OHP GSL signaling, we analyzed mutants of two key autophagic (ATG) components, *atg2-1* (18) and *atg5-1* (58). ATG2 is part of the ATG9 cycling system that is essential for autophagosome formation (19, 59, 60). ATG9-containing vesicles are a suggested membrane source for the autophagosome, and vesicles containing ATG9 are cycled to-and-from the phagophore via the ATG9 cycling system (19, 60). ATG5, is part of the dual ubiquitin-like conjugation systems responsible for ATG8 lipidation (19, 60, 61). There are nine *ATG8* paralogues in Arabidopsis (60) and together with the single copy of *ATG5*, they are essential for autphagosome initiation, expansion, closure, and vacuolar fusion (19, 60). After the first conjugation system has conjugated ATG8 to an E2-like enzyme, the E3 ligase-like activity of the second ATG5-containing system enables ATG8 lipidation at the autophagic membrane (19, 62, 63). We found that *atg5-1* enhanced 3OHP GSL responsiveness (Figure 8A-B) while *atg2-1* had a wild type response (Figure 8C-D). One possible explanation for this difference between the two mutants is that, apart from macro-autophagy, plants also have micro-autophagy (60), a process that, in animal systems, has been shown to be negatively regulated by TOR (64). Micro-autophagy does not involve *de novo* assembly of autophagosomes, and ATG5 has been shown to be involved in several forms of micro-autophagy whereas the role of ATG2 is more elusive and may not be required (64). Thus, the elevated 3OHP GSL response in the *atg5-1* mutant supports the hypothesis that 3OHP GSL signaling proceeds through the TOR pathway, but also suggests that this response requires parts of the autophagic machinery as it was not observed for *atg2-1*.

**Figure 8.**
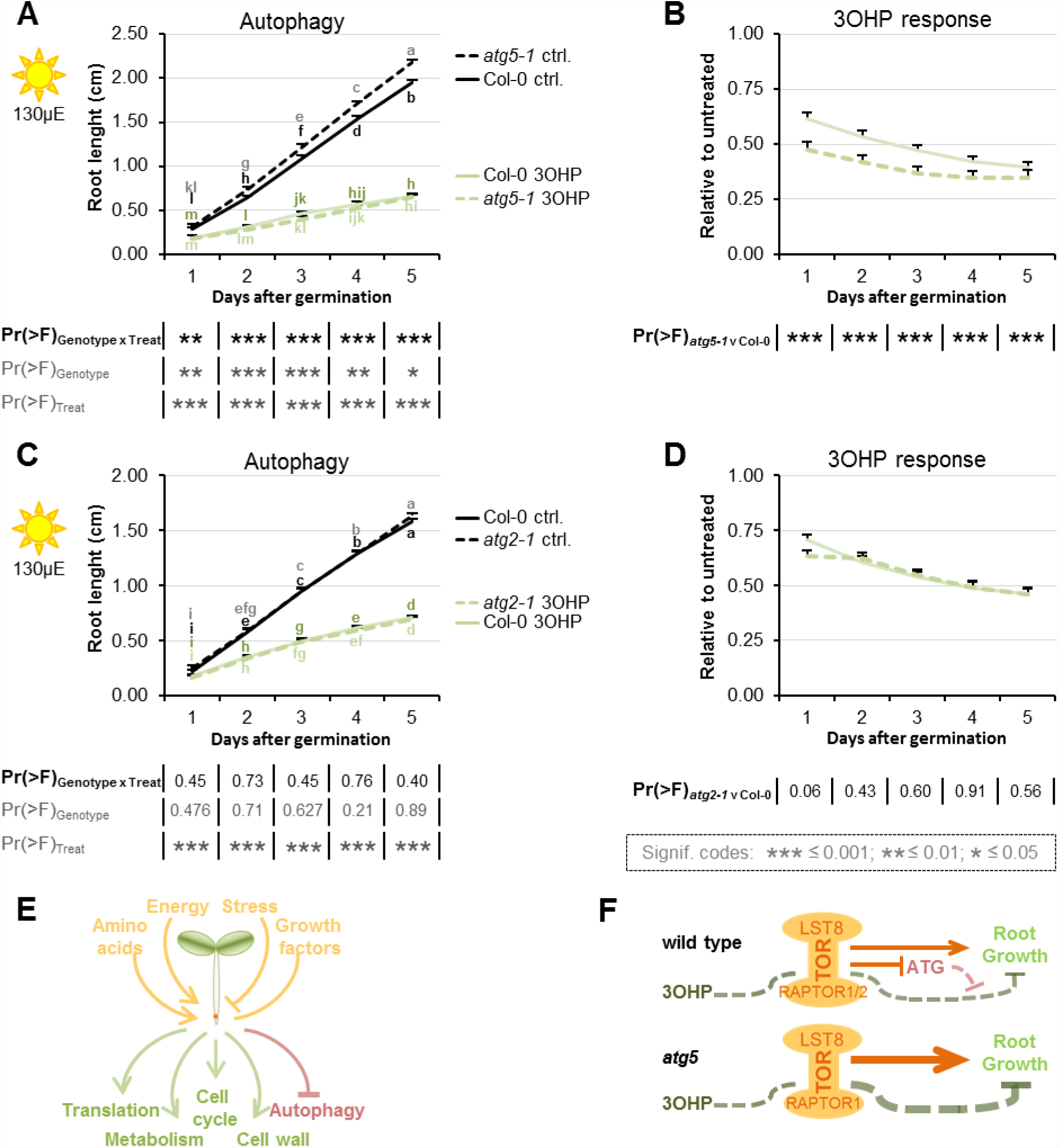
Blocking autophagosome elongation amplifies the 3OHP response. **A** Root growth for *atg5-1* and wildtype Col-0 seedlings grown on MS medium supplemented with or without 5μM 3OHP. Multi-factorial ANOVA was used to test the impact of Genotype (Col-0 v *atg5-1*), Treatment (Control v 3OHP) and their interaction on root length. All experiments were combined in the model and experiment treated as a random effect. The ANOVA results from each day are presented in the table. **B** Root lengths in response to 3OHP (from A) displayed at each time point as relative to untreated. Results are least squared means ± SE over two independent experimental replicates with each experiment having an average of 21 replicates per condition (n=31-52). **C** Root growth for *atg2-1* and wildtype Col-0 seedlings grown on MS medium supplemented with or without 5μM 3OHP. Multi-factorial ANOVA was used to test the impact of Genotype (Col-0 v *atg5-1*), Treatment (Control v 3OHP) and their interaction on root length. All experiments were combined in the model and experiment treated as a random effect. The ANOVA results from each day are presented in the table. **D** Root lengths in response to 3OHP treatment (from C) displayed at each time point as relative to untreated. Results are least squared means ± SE over two independent experimental replicates with each experiment having an average of 26 replicates per condition (n=36-66). **E** The TOR complex (TORC), is affected by several upstream input, leading to activation or repression of several downstream pathways. **F** Schematic model; sucrose activates TORC, leading to root growth. 3OHP represses root growth through interaction with TORC. Autophagy pathways via ATG5 negatively affect 3OHP response.

## Discussion

In this study we describe a novel signaling capacity associated with 3OHP GSL, a defense metabolite present in the Brassicaceae Arabidopsis, and provide evidence that the linked signal proceeds via the TOR pathway. Application of exogenous 3OHP GSL caused reversible root meristem inhibition by morphological reprogramming of the root zones, i.e. dramatically reduced the root meristem size and limited root cell elongation (Figure 4). This response occurred at levels within the endogenous range and there was no evidence of cell death in any treated root, suggesting that this is not a toxicity response (Figure 1). Additionally, these morphological responses were specific to 3OHP GSL and not caused by any structurally or biosynthetically related GSL, suggesting that these responses were not because of generic properties shared by GSLs (Figure 2). Exposing a wide phylogenetic array of plants, including lineages that have never produced GSLs, to 3OHP GSL showed that application of this compound can inhibit growth broadly across the plant kingdom as well as in yeast (Figure 3, Figure 3 –figure supplement 1). This suggests conservation of the downstream signaling pathway across these diverse plant lineages. Equally, if the signaling compound is not 3OHP GSL itself, but a derivative, then the required biosynthetic processes must be conserved. This conservation largely rules out the specific GSL activation pathway controlled by Brassicales specific thioglucosidases, myrosinases (65-67). The phylogenetic conservation of the 3OHP GSL response led us to search for a target pathway controlling growth and development that would be evolutionary well conserved between the tested species.

By comparing the root phenotype identified with 3OHP GSL application to the published literature, we hypothesized that 3OHP GSL treatment may affect TORC, a key primary metabolic sensor that controls growth and development, and is conserved back to the last common eukaryotic ancestor (2). Active site TOR inhibitors inhibit root growth in numerous plant species similar to 3OHP GSL application (56), supporting the hypothesis that 3OHP GSL may function via TORC. A model with 3OHP GSL affecting TORC would explain how 3OHP GSL can alter root development across the plant kingdom (Figure 3). Mechanistic support for this hypothesis came from a number of avenues. First, 3OHP GSL can block the TOR-mediated sugar activation of arrested meristems (Figure 6). Second, the TORox mutant intensifies 3OHP GSL linked signaling (Figure 5), and correspondingly loss-of-function mutants of the substrate binding TORC component *raptor2* diminish the 3OHP GSL effect (Figure 5 –figure supplement 3). Additionally, there are clear phenotypic overlaps between the root phenotypes induced by known TOR inhibitors and 3OHP GSL, e.g. root inhibition, inhibition of cell elongation, and notably the dramatic reduction of the meristem sizes (Figure 7). Critically, 3OHP GSL and known TOR inhibitors were antagonistic for a number of phenotypes. In pharmacology, the outcomes of a drug combination can either be antagonistic, additive or synergistic, depending on whether the effect is less than, equal to, or greater than the sum of the effects of the two drugs (57). Antagonistic interactions, as observed with 3OHP GSL and AZD, can occur if two drugs exhibit mutual interference against the same target site, or if their targets converge on the same regulatory hub (57). Together, these lines of evidence suggest that 3OHP GSL targets the TOR pathway to alter root meristem development within Arabidopsis and potentially other plant species.

Extending the analysis to pathways downstream of TORC, showed that loss of ATG5, a vital component of the autophagic machinery (16, 17), intensifies the 3OHP GSL response (Figure 8A-DB). This supports the hypothesis that 3OHP GSL signaling proceeds through the TOR complex, but also suggest that this signal requires parts of the autophagic machinery. Loss of another autophagic component, ATG2, did not influence the 3OHP response (Figure 8C-D). Together, this raises the possibility that 3OHP GSL responses involve predominantly a micro-autophagy pathway, which is ATG5- but may not be ATG2-dependent, rather than the macro-autophagy pathway that depends upon both genes (60) (64). Micro-autophagy removes captured cytoplasmic components directly at the site of the vacuole via tonoplast invagination. The cargo to be degraded ranges from non-selective fractions of the cytoplasm to entire organelles, dependent on the type of micro-autophagy. The two ubiquitin-like conjugation systems, and thereby ATG5, have been shown to be involved in several forms of micro-autophagy, such as starvation-induced, non-selective, and glucose-induced selective autophagy (64). Interestingly, micro-autophagy involves vacuolar movement of cargo, and the vacuole is considered the main storage site for glucosinolates [51]. Thus, ATG5 may be responsible for enabling the movement of exogenously applied 3OHP GSL out of the cytoplasm where it could interact with the TORC pathway and into the vacuole. This would decrease the concentration of the 3OHP GSL signal and could explain why the *atg5-1* mutant is more sensitive to 3OHP GSL. Further work is required to test if ATG5 is functioning to attenuate the 3OHP GSL signal.

A conundrum for defense signaling compounds to affect growth is the evolutionary age discrepancy; defense metabolites are typically evolutionarily very young, as they are often species or taxa specific, while growth regulatory pathways are highly conserved across broad sets of plant taxa. This raises the question of which mechanism(s) may allow this connection between young metabolites and old regulatory pathways. This suggests that plants may sense young metabolites using evolutionarily old signaling pathways. Similar evidence is coming from other secondary metabolite systems suggesting that this may be a general phenomenon. For example, an indolic GSL activation product can interact with the conserved *TIR1* auxin receptor to alter auxin sensitivity within Arabidopsis (23). Similarly, an unknown phenolic metabolite appears to affect regulation of growth and development by influencing the Mediator complex that is conserved across all eukaryotes (24-26); and the plant polyphenol resveratrol directly inhibits the mammalian TOR to induce autophagy (68). Thus, young plant metabolites can influence evolutionarily conserved pathways. Interestingly, this strongly resembles the action of virulence-associated metabolites within plant pathogens. Pseudomonad bacteria produce the evolutionarily young coronatine that alters the plant defense response by interacting with the conserved JA-Ile receptor *COI1* (69). In plant/pathogen interactions, this ability of pathogen-derived metabolites to alter plant defense signaling is evolutionarily beneficial because it boosts the pathogens virulence *in planta*. It is less clear if this selective pressure model also applies to plant defense compounds that interact with endogenous signaling pathways. Following the plant/pathogen derived model, such plant defense metabolites might have been co-selected on the ability to affect the biotic attacker and simultaneously provide information to the plant. However, these examples may simply be serendipitous cases, where the defense metabolite happened to interact with a pathway. Testing these hypotheses would require a broad survey to assess how many plant metabolites can affect signaling within the plant. This survey would also be required to assess how frequently defense metabolites may play signaling roles within plants and what the targeted pathways are.

Within this report, we provided evidence that 3OHP GSL, or derived compounds, appears to function as a natural endogenous TORC inhibitor that can work across plant lineages. This creates a link whereby the plant’s endogenous defense metabolism can simultaneously coordinate with growth. Such a built-in signaling capacity would allow coordination between development and defense, as the plant could use the defense compound itself as a measure of the local progress of any defense response and readjust development and defense to optimize against the preeminent threat. Future work is required to identify the specific molecular interaction that allows this communication to occur, this will help to illuminate how and why plants measure their own defense metabolism to coordinate available resources more broadly with growth. Future work might also ascertain whether there is a broader class of plant produced TOR inhibitors. If this is true, they might be highly useful in understanding TOR function across kingdoms of life and possibly to reveal significant aspects of this universally conserved pathway that may have gone unnoticed in other eukaryotic models.

## Materials and methods

### Plant Materials

The genetic background for the Arabidopsis (*Arabidopsis thaliana*) mutants and transgenic lines described in this study is the Col-0 accession. The following lines were described previously: *myb28-1 myb29-1* (70)*, atg2-1* (18)*, atg5-1* (58)*, raptor1-1* (54, 55), *raptor1-2* (54)*, raptor2-2* (54), *raptor2-1* (54, 55), and the *TORox* lines *G548, G166, S784*, and *S7817* (53). All genotypes were obtained and validated both genetically and phenotypically as homozygous for the correct allele.

### Plant growth media and *in vitro* root growth assays

Seeds were vapour sterilized for 2-3 hours, by exposure to a solution of 100 mL household bleach (Klorin Original, Colgate-Palmolive A/S) mixed with 5 mL hydrochloric acid (12M), and ventilated for 30 min to one hour. After plating, on ½ strength Murashige and Skoog (MS) medium (2.2 g/l MS+vitamins) (Duchefa) with 1% (w/v) sucrose (Nordic Sugar), and 0.8% (w/v) micro agar (Duchefa), pH adjusted to 5.8), the seeds were stratified for two days in the dark at 4°C. For root length assays at normal light (115-130μE) Arabidopsis seedlings were grown vertically at 22°C day 20°C night under a 16-h photoperiod and 80% humidity (long day). For meristem reactivation assays plants were grown as described in (12), except that in our conditions we needed to go to 25μE to obtain meristem inhibition. Daily root lengths were manually marked (from day 3) with a permanent marker pen on the backside of the plate. After photography of 7-d-old seedlings the root growth was quantified using the ImageJ software(71). The least square means (lsmeans) for the genotypes in response to different treatments were calculated across experiments (in R, see statistics), and plotted in excel.

### *in vitro* root growth assays for the species (seed plating and growth conditions)

To test 3OHP GSL perception in other plant orders, seeds were obtained as listed in table S1. Except for *Solanum lycopesicum*, here *San Marzano* tomatoes were bought in a local supermarket and the seeds were harvested, fermented and dried. All seeds were vapour sterilized for three hours (as above). Before plating, and *Lotus japonicus MG20* (Lotus) were emerged in water and kept at 4°C for 1-2 weeks. Seeds were plated on vertical ½MS plates as specified in table S1, stratified for four days in the dark at 4°C before being transferred to a long day growth chamber). Root growth was measured approximately every 24 hours (as described above).

### Yeast strain, media, and growth conditions

The yeast strain, NMY51 with pOST1-NubI and pDHB1-LargeT ((72, 73); DUALsystem Biotech), was grown in liquid YPD media (2% w/v bactopeptone (Duchefa Biochemie), 1% w/v yeast extract (Becton, Dickinson and Company), 2% w/v glucose) with or without added GSLs, at 30°C and 150 rpm shaking.

### Yeast growth assay

On day one; a 5 ml overnight culture was started from cryostock. Day two; four new 4ml cultures were inoculated with 1 ml overnight culture, and grown overnight. On day three; an OD600 0.4 and a 0.04 dilution was prepared from each of the four cultures. 500 μl of each of the four cultures, at both dilutions, were transferred to a 96-well culture plate containing 500 μl YPD liquid media with 3OHP GSL or Allyl GSL, to final OD600 0.2 and GSL concentrations of 50, 10, 5, 1 and 0 μM. The yeast growth was measured at 0, 4, 6, 8, 24 and 48 hours. For each growth measurement 100 μl culture was transferred to a 96-well Elisa-plate together with three wells of YPD liquid media for standardization. Growth was measured with a SpectraMAX 190 (Molecular Devices) and SoftMax^®^ Pro 6.2.2 software. Growth rates and statistical analysis was calculated using the R software. The linear growth range was determined, and a linear regression using the lm() function in R was carried out to determine OD600 increase per hour (slope) and the yeast doubling time was calculated.

### Glucosinolate Analysis

Glucosinolates were extracted from whole plant tissue of adult plants (for 3OHP GSL extraction), or from or 10-d-old seedlings (3OHP GSL uptake) (44, 74, 75), and desulfo-glucosinolates were analysed by LC-MS/TQ as desulfo-GSLs as described in (76).

### Statistics

The R software with the R studio interface was used for statistical analysis (77, 78). Significance was tested using the Anova function (aov), lsmeans were obtained using the ‘lsmeans’ package (version 2.17) (79). The letter groupings (Tukey’s HSD Test) were obtained using the ‘agricolae’ package (version 1.2-3) (80).

### Confocal Microscopy

To examine the root tip zones, we used confocal laser-scanning microscopy of 4-d-old seedlings grown vertically with or without treatment (with 3OHP GSL and/or various inhibitors). Samples were mounted on microscopy slides in propidium iodide solution (40μM, Sigma) and incubated for 15 minutes. Confocal laser scanning microscopy was carried out on a Leica SP5-X confocal microscope equipped with a HC PL FLUOTAR 10 DRY (0.3 numerical aperture, 10X magnification) or a HCX lambda blue PL APO 320 objective (0.7 numerical aperture, 20X magnification) for close-up pictures of the meristem. To visualize the cell walls of individual cells the propidium iodide stain was excited at 514 nm and emission was collected at 600 nm to 680 nm. To determine the size of the meristems the confocal pictures we manually inspected and the meristematic cells marked and counted (the meristem region is defined as in (1, 81)). To measure the distance from the root tip to the point of first root hair emergence we used ImageJ (71).

### Chemicals

The AZD8055 (82), Torin2 (83), KU-63794 (84), and WYE-132 (85) were purchased from Selleckchem. PF-4708671 (86) and allyl/sinigrin were purchased from Sigma-Aldrich. 4MSB and 3MSP GSLs were purchased at C2 Bioengineering. But-3-enyl GSL was purified from *Brassica rapa* seeds while 3OHP GSL was purified from the aerial parts of 4-5weeks old greenhouse-grown plants of the Arabidopsis accession Landsberg *erecta* (75, 76). The concentration of 3OHP and but-3-enyl GSL was determined by LC-MS/TQ as desulfo-GSLs. All inhibitors were dissolved in DMSO and stored as 10mM stocks at –20 °C. For allyl, 3MSP, and 4MSB ~100mM GSL stocks were made with H2O and the concentration of GSLs within these stocks was determined by LC-MS/TQ (see above).

## Funding

Funding for this work was provided by the Danish National Research Foundation (DNRF99) grant to DJK and MB, the NSF award IOS 13391205 and MCB 1330337 to DJK, and the USDA National Institute of Food and Agriculture, Hatch project number CA-D-PLS-7033-H to DJK.

## Acknowledgements

We thank the excellent technical assistance of the PLEN Greenhouse staff, and the DynaMo student helpers. We thank Dr. Svend Roesen Madsen for providing Camelina, and rape seeds, and Dr. Camilla Knudsen Baden for giving us lotus seeds.

## Conflict of Interest Statement

The authors declare that the research was conducted in the absence of any commercial or financial relationships that could be construed as a potential conflict of interest.

---

**Figure 1 –figure supplement 1.**
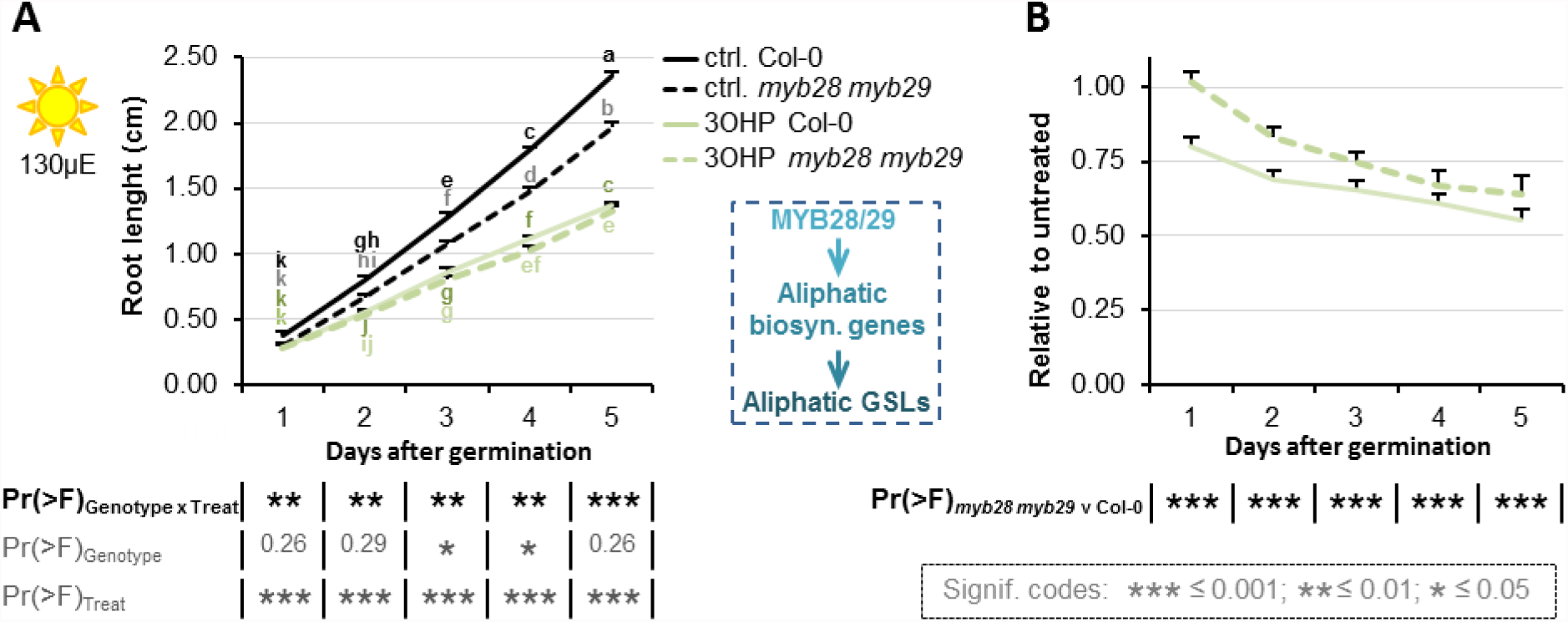
Root inhibition is affected by endogenous 3OHP/GSLs levels. Results are least squared means ± SE over three independent experimental replicates with each experiment having an average of ten replicates per condition (n=8-39). A Root growth for seedlings grown on MS medium supplemented with or without 5μM 3OHP. Multi-factorial ANOVA was used to test the impact of Genotype (Col-0 v myb28myb29), Treatment (Control v 3OHP) and their interaction on root length. The ANOVA results from each day are presented in the table. B Root lengths in response to 3OHP (from A) displayed at each time point as relative to the untreated

**Figure 3 –figure supplement 1.**
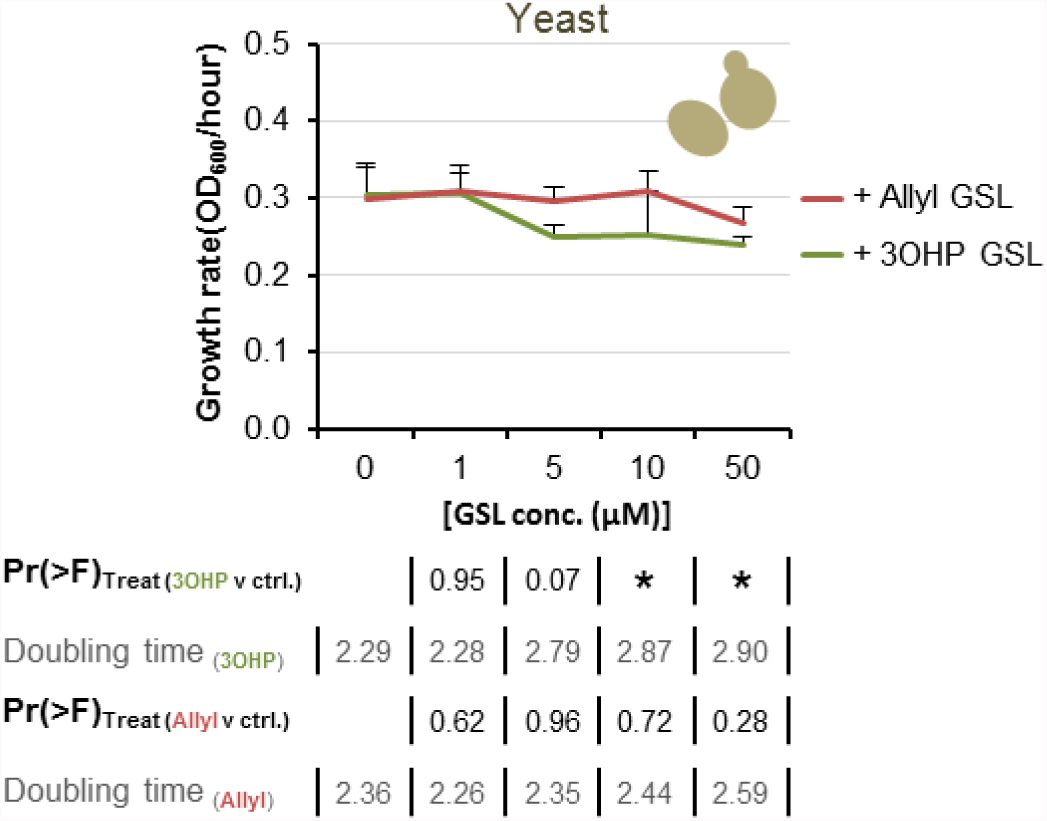
Yeast response to 3OHP suggests a conserved target throughout eukaryotes. Yeast growth in YPD media supplemented with none or increasing levels of 3OHP or Allyl. The hourly OD600 increase is plotted against each concentration of either Allyl or 3OHP. The least squared means ± SE over four replicates are presented (n=4). ANOVA was utilized to test for a significant effect of GLS treatment individually for each concentration of 3OHP and Allyl.

**Figure 5 –figure supplement 1.**
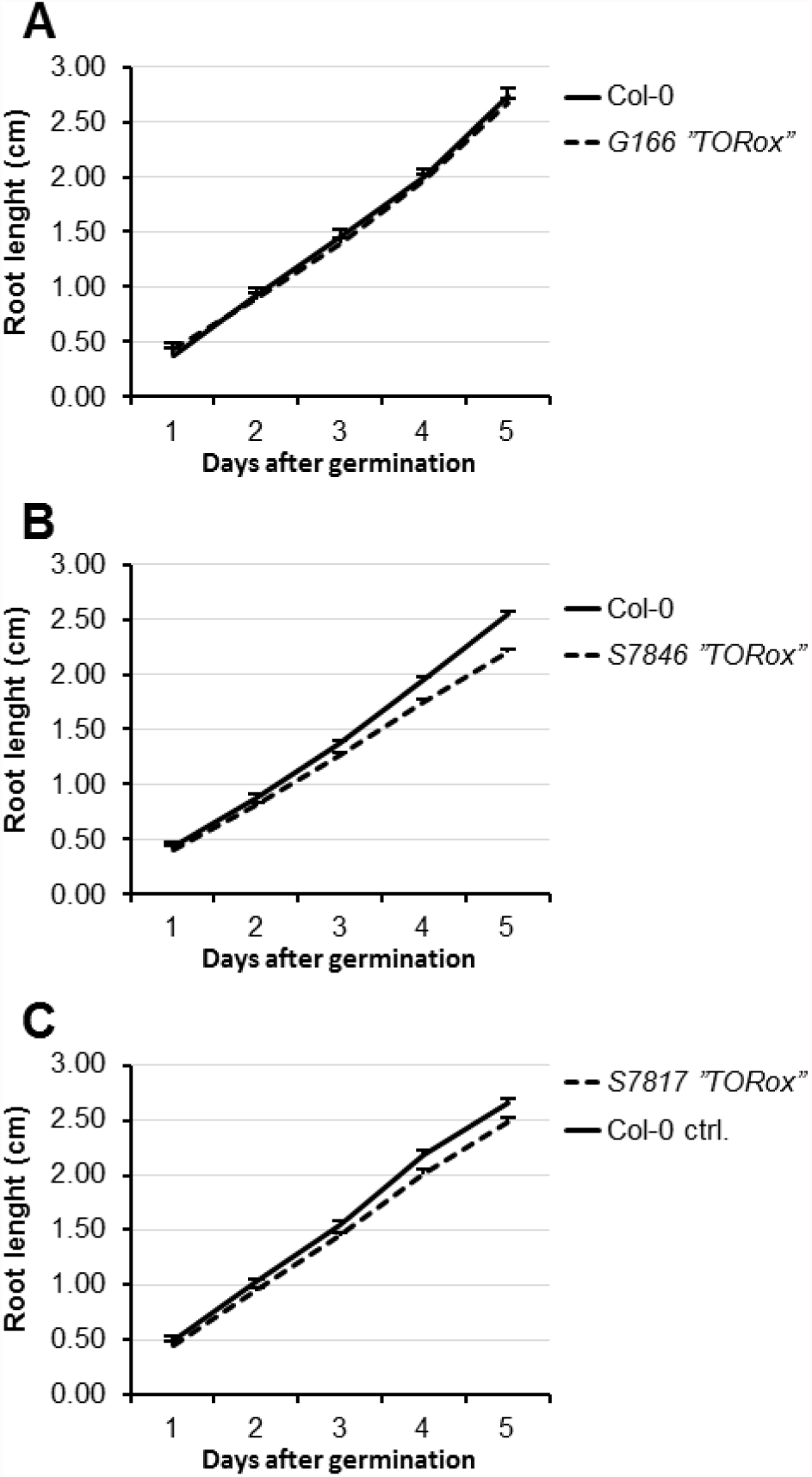
Published TORox lines that did not display the TORox phenotype under our conditions. Multi-factorial ANOVA was used to test the impact of Genotype (Col-0 v specific TORox lines) on root length. All experiments were combined in the model and experiment treated as a random effect. There were no significant differences found. **A** Root growth for the published TORox line G166 and wildtype Col-0 seedlings grown on MS medium supplemented with or without 5μM 3OHP. Results are least squared means ± SE (n=8-16). **B** Root growth for the published TORox line S7846 and wildtype Col-0 seedlings grown on MS medium supplemented with or without 5μM 3OHP. Results are least squared means ± SE ns across three biological repeats (n=35-45). **C** Root growth for the published TORox line S7817 and wildtype Col-0 seedlings grown on MS medium supplemented with or without 5μM 3OHP Results are least squared means ± SE (n=10-24).

**Figure 5 –figure supplement 2.**
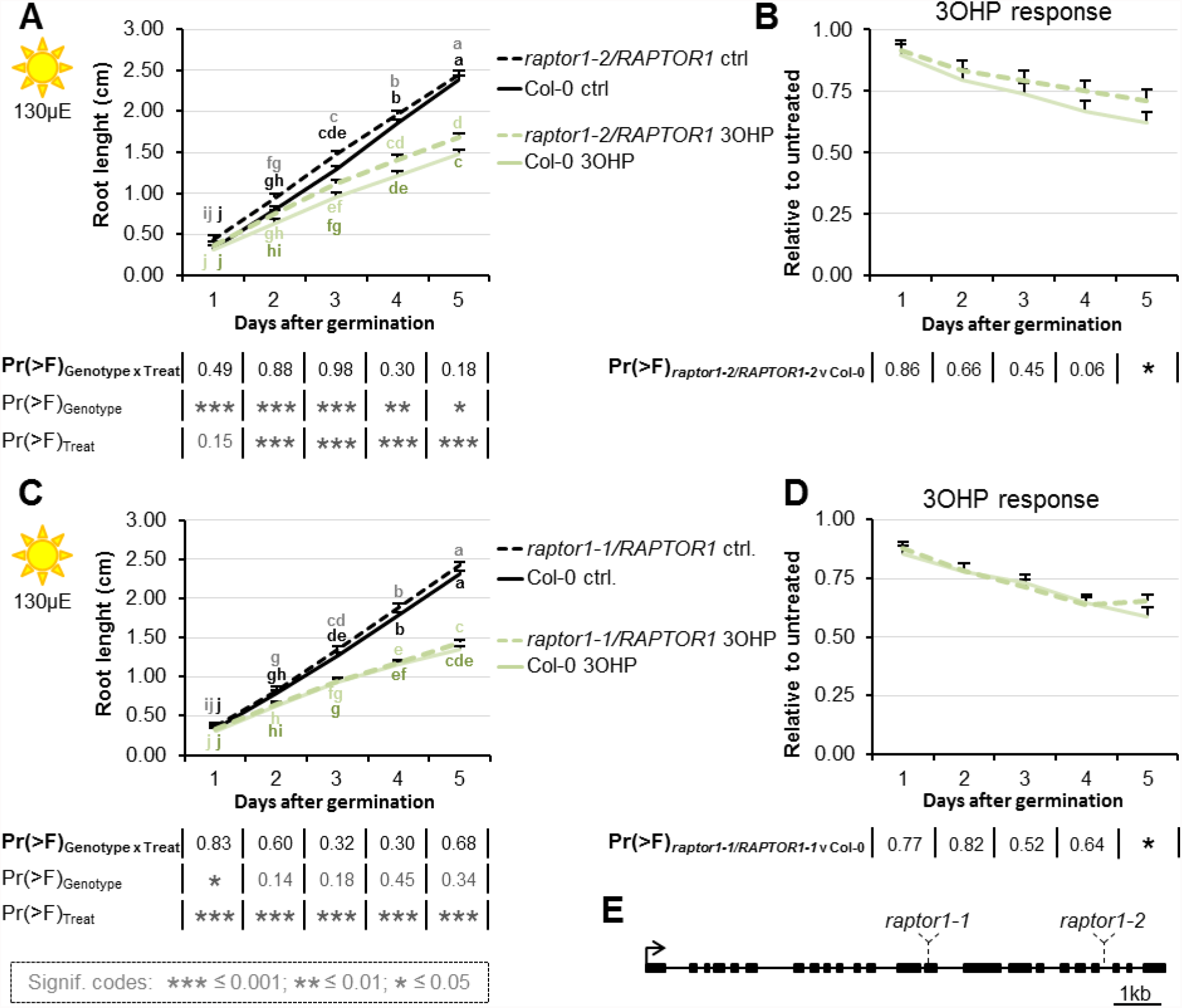
*RAPTOR1* haplo-insufficiency does not affect 3OHP response. **A** Root growth for heterozygous *raptor1-2* and wildtype Col-0 seedlings grown on MS medium supplemented with or without 5μM 3OHP. Multi-factorial ANOVA was used to test the impact of Genotype (Col-0 v *raptor1-2*), Treatment (Control v 3OHP) and their interaction on root length. All experiments were combined in the model and experiment treated as a random effect. The ANOVA results from each day are presented in the table. **B** Root lengths in response to 3OHP (from A) displayed at each time point as relative to untreated. Results are least squared means ± SE over three independent experimental replicates with each experiment having an average of six replicates per condition (n=16-19). **C** Root growth for heterozygous *raptor1-2* and wildtype Col-0 seedlings grown on MS medium supplemented with or without 5μM 3OHP. Multi-factorial ANOVA was used to test the impact of Genotype (Col-0 v *raptor1-1*), Treatment (Control v 3OHP) and their interaction on root length. All experiments were combined in the model and experiment treated as a random effect. The ANOVA results from each day are presented in the table. **D** Root lengths in response to 3OHP (from C) displayed at each time point as relative to untreated. Results are least squared means ± SE over three independent experimental replicates with each experiment having an average of seven replicates per condition (n=16-24). **E** Gene structure and T-DNA insertion sites for *RAPTOR1*.

**Figure 5 –figure supplement 3.**
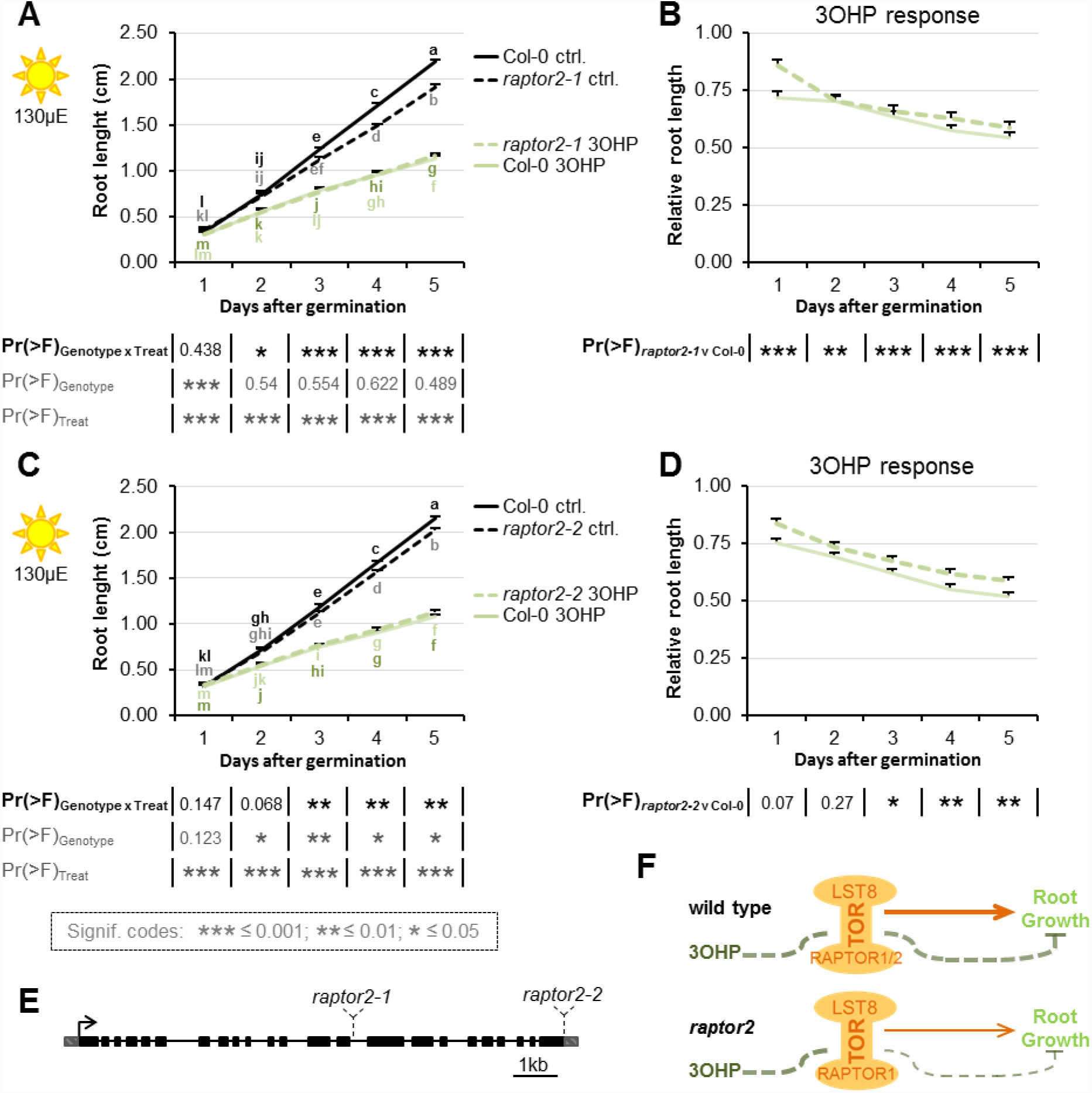
Loss of one of the two substrate-binding TORC-subunits affect 3OHP response. **A** Root growth for *raptor2-1* and wildtype Col-0 seedlings grown on MS medium supplemented with or without 5μM 3OHP. Multi-factorial ANOVA was used to test the impact of Genotype (Col-0 v *raptor2-1*), Treatment (Control v 3OHP) and their interaction on root length. All experiments were combined in the model and experiment treated as a random effect. The ANOVA results from each day are presented in the table. **B** Root lengths in response to 3OHP (from A) displayed at each time point as relative to untreated. Results are least squared means ± SE over four independent experimental replicates with each experiment having an average of thirteen replicates per condition (n=36-68). **C** Root growth for *raptor2-2* and wildtype Col-0 seedlings grown on MS medium supplemented with or without 5μM 3OHP. Multi-factorial ANOVA was used to test the impact of Genotype (Col-0 v *raptor2-1*), Treatment (Control v 3OHP) and their interaction on root length. All experiments were combined in the model and experiment treated as a random effect. The ANOVA results from each day are presented in the table. **D** Root lengths in response to 3OHP (from C) displayed at each time point as relative to untreated. Results are least squared means ± SE over three independent experimental replicates with each experiment having an average of nineteen replicates per condition (n=44-70). **E** Gene structure and T-DNA insertion sites for *RAPTOR2*. **F** Schematic model; loss of one of the substrate-binding subunits RAPTOR2 decreases growth, and the relative 3OHP response.

**Figure 7 –figure supplement 1.**
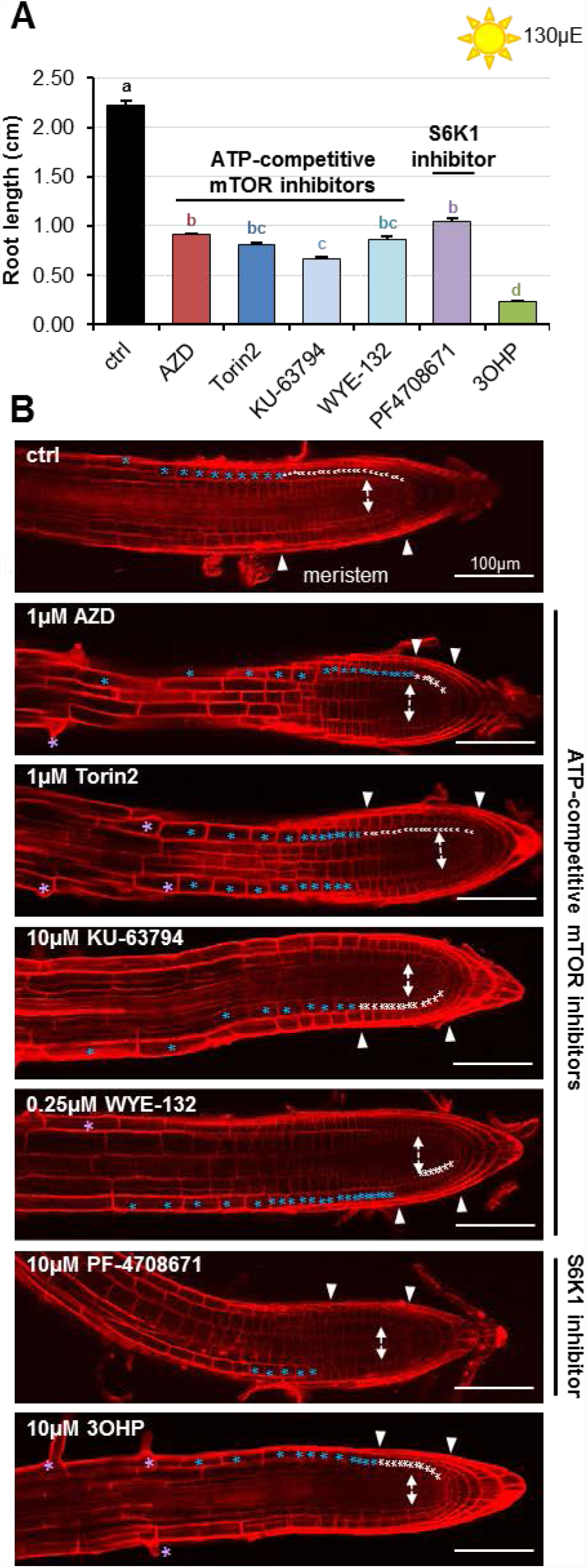
**A**. Root lengths of 7-d-old Col-0 wildtype seedlings grown on MS medium with sucrose ± the indicated mTOR or S6K inhibitors. Results are averages ± SE (n=8-41). **B** Confocal images of 4–d-old propidium iodide stained seedlings. Meristemal cells are marked with white asterisks, elongated cells with blue, and cells belonging to the differentiation zone are marked with purple asterisks. Arrows indicate approximate meristem sizes.

**Table S1.**
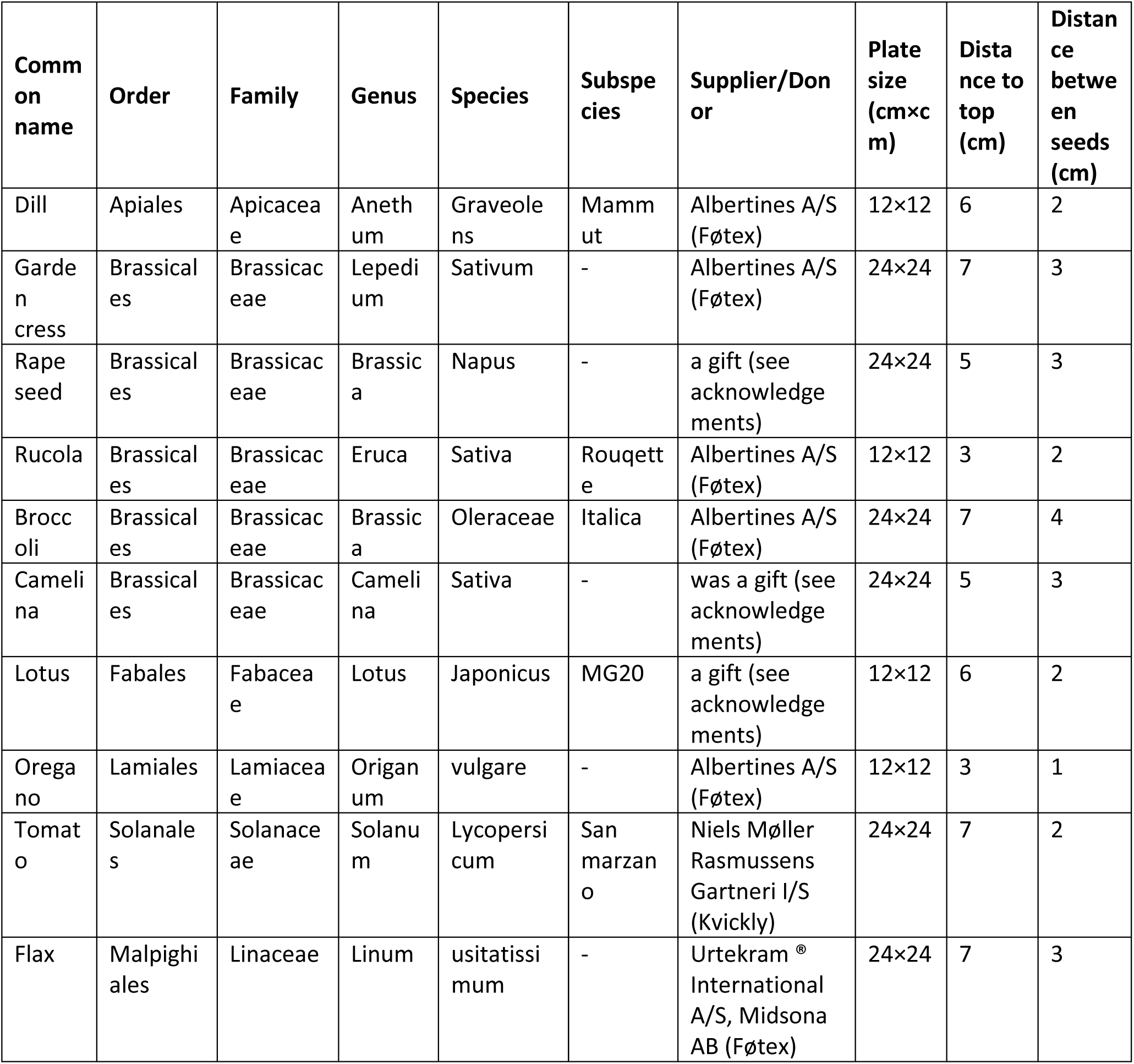
Sup. Table 1. Plant seeds used for *in vitro* root growth assays for the various plant species. The, source, common name, order, family, genus, species, and subspecies seeds are listed for the used plant species. As well as the plate size (cm×cm), and plating distance used for the individual response assays.

